# Age-informed, attention-based weakly supervised learning for neuropathological image assessment

**DOI:** 10.1101/2025.05.16.654510

**Authors:** Shuying Li, Maxwell Malamut, Ann McKee, Jonathan D. Cherry, Lei Tian

## Abstract

Chronic Traumatic Encephalopathy (CTE) and other neurodegenerative disorders (NDs) pose diagnostic challenges due to their diffuse and subtle pathological changes. Traditional diagnostic methods relying on manual histopathological slide inspection are labor-intensive and prone to variability, often missing subtle structural alterations. This study introduces an age-informed, attention-based multiple instance learning (MIL) pipeline to predict AT8 density, a key marker of p-tau aggregation in CTE. Using Luxol Fast Blue and Hematoxylin & Eosin (LH&E) stained images, our model identifies critical pathological regions and generates interpretable attention maps highlighting structural changes linked to tau pathology. Incorporating patient age enhances predictive accuracy and contextual understanding, addressing aging’s confounding effects. We also develop quantitative evaluation procedures for foundation models (FMs), assessing attention map smoothness, faithfulness, and robustness to perturbations like stain variability and noise. These benchmarks facilitate informed FM selection and optimization for neuropathological tasks. By enabling scalable, automated whole-slide image (WSI) analysis, our approach advances digital neuropathology, supporting earlier and more precise ND diagnoses and uncovering subtle markers with potential applications in clinical imaging.

## 1 Introduction

Neurodegenerative disorders (NDs) such as Chronic Traumatic Encephalopathy (CTE) pose a substantial burden on individuals and society due to their significant and enduring impacts on cognitive function and mental health [1, 2], underscoring the urgent need for accurate and comprehensive diagnostic methods. Currently, the diagnostic criteria for CTE heavily depend on the manual inspection of histopathological images [3, 4], a process that is both labor-intensive and subject to considerable intra- and inter-observer variability. Diagnostic workflows traditionally rely on evaluating known markers, such as hyperphosphorylated tau (p-tau) [3–5], with changes also observed in Luxol Fast Blue and Hematoxylin and Eosin (LH&E) stained slides [6]. While structural alterations observed in LH&E slides, such as cortical atrophy, white matter rarefaction, and perivascular pigment-laden macrophages, have been associated with CTE [6], there may be subtle changes currently overlooked, underscoring the need for more scalable, accurate, and reproducible assessment methods.

Recent advancements in digital pathology and deep learning hold significant potential for addressing the challenges in neuropathology by enabling automated analyses of neuropathological images. Foundation models (FMs) [7–15]— large, pre-trained neural networks providing generalizable feature representations—and multiple instance learning (MIL) [16, 17]— a weakly supervised learning approach that assigns slide-level rather than pixel-level labels — stand out as particularly promising strategies for analyzing whole-slide images (WSIs). These methods can reduce reliance on labor-intensive manual annotations while improving diagnostic precision.

However, adapting these computational approaches to neuropathology presents unique obstacles. Unlike cancer pathology, which often involves localized lesions, neurodegenerative diseases like CTE exhibit diffuse and subtle changes distributed across multiple brain regions [6, 18]. This lack of clearly delineated regions of interest (ROIs) complicates the evaluation of attention maps. Moreover, the brain’s intricate architecture, composed of distinct regions and cellular layers, means pathological changes can vary significantly between regions and layers [3, 4, 6, 18]. Finally, the confounding effects of aging introduce additional variability [18] necessitating the integration of age into computational models.

In this work, we present an age-informed attention-based MIL pipeline for neuropathology. We use the prediction of AT8 density, a key marker for p-tau aggregation in CTE [18], as a case study to demonstrate the pipeline’s capabilities for providing neuropathological insights. Leveraging commonly available LH&E staining techniques, this pipeline predicts AT8 density and generates attention maps that identify critical pathological regions with structural changes associated with known patterns of tau pathology in CTE. The inclusion of age as an informative feature in our model enables more accurate predictions and better contextual understanding of the observed tau pathology. Additionally, we offer comprehensive quantitative procedures to evaluate a range of existing FMs, enabling the informed selection of FMs most appropriate to neuropathological applications and guiding the future development of neuropathology-specific FMs. We assess the smoothness and faithfulness of the attention maps, even in the absence of annotated ROIs. Furthermore, we provide quantitative measures to evaluate the model’s robustness to common perturbations in neuropathology, including stain variations and random noise. By doing so, our approach advances the application of deep learning in digital neuropathology, improving both the diagnosis of NDs and the identification of novel disease markers. This scalable solution enables large-scale analysis of neuropathological WSIs, offering new insights into the pathology of NDs such as CTE, and addressing key challenges in neuropathology.

## 2 Related work

### 2.1 Foundation Models (FMs) in Digital Pathology

Analyzing large WSIs typically involves compressing their rich content into a lower-dimensional space using pre-trained large-scale FMs. These models not only enhance computational efficiency and analysis accuracy but also represent a major shift in data-intensive science methodologies. They are pivotal for automating image analysis and uncovering pathological markers and disease subtypes that are challenging or impossible for human experts to detect. By standardizing and scaling analyses across thousands of WSIs, these models contribute to more robust and reproducible findings, thereby mitigating the subjectivity in manual interpretations.

Several FMs for digital pathology in oncology have been developed recently [7–15]. The pioneering model CTransPath utilized a Swin transformer architecture, trained with MoCoV3 on 15 million tiles from 32,000 WSIs [9]. More recent studies have increasingly adopted DINOv2 as preferred foundational architectures in digital pathology[10, 11, 15]. The training datasets for these models have grown to 0.1 billion to more than 1 billion tiles, derived from 0.1 million to over 1 million WSIs [7, 10, 11]. Notably, UNI [11] was trained on one of the largest histology slide datasets, comprising over 100 million tissue patches from 100,426 diagnostic H&E WSIs spanning 20 major tissue types. It demonstrated robust performance across a range of downstream tasks in oncology, including cell type segmentation, ROI-level classification, and slide-level classification. Prov-GigaPath[10], trained on an extensive dataset of 1.38 billion tiles extracted from 171,189 H&E-stained and immunohistochemistry pathology slides representing 31 major tissue types from more than 30,000 patients, offers both patch-level and whole-slide-level representations. The Virchow FM, trained on data from approximately 100,000 patients and around 1.5 million WSIs, has achieved superior performance compared to previous models [7]. These FMs have primarily been evaluated based on their performance in downstream tasks. A significant gap remains in the development of comprehensive evaluation strategies for selecting FMs, which is essential for ensuring robust model performance in neuropathological contexts.

### 2.2 Multiple instance Learning (MIL)

MIL[19] is a weakly supervised learning approach that can effectively train deep neural networks when fully annotated data is unavailable. It can classify WSIs based on a single slide-level diagnosis (e.g., a “ground-truth” label), rather than relying on pixel-level annotations (e.g., segmented regions of interest). Thus, MIL is particularly beneficial in digital pathology, where labels are frequently provided for WSIs but are less common for smaller patches, regions, or pixels. MIL treats the whole image as a “bag” of data, with multiple objects called “instances” belonging to potentially different implicit classes. Applied to WSIs, most MIL-based approaches divide the image into patches of manageable size for computational processing and treat each patch as an instance in the WSI bag.

A notable enhancement in MIL was introduced by Ilse et al. [16]who incorporated an attention mechanism that automatically learns to suppress instances unrelated to the target class. Furthermore, the attention mechanism yields a weight for each instance that quantifies its contribution to the overall classification and generates attention maps that highlight ROIs, allowing models to provide interpretable insights into their focus without requiring pixel-level expert annotations [16]. These attention maps enable efficient and scalable visualization of critical regions by directly leveraging slide-level labels, facilitating model evaluation and feature detection. Lu et al. [20] extended this approach by adding a clustering constraint for digital pathology applications. This method uses the separation between instances with the highest and lowest attention weights within a bag as a constraint to refine the classification process. The resulting method, CLAM, has been shown to outperform traditional MIL methods in cancer detection and subtyping. In oncology, MIL-based methods have achieved success by efficiently processing WSIs at scale for tasks such as tumor classification, disease staging and subtyping, and predicting survival outcomes [17, 20–24].

### 2.3 Deep Learning-based Integration of Patient Metadata with Medical Imaging

Integrating clinical data such as age and sex into medical images has been shown to enhance the performance of deep-learning models. Various integration strategies have been explored, including concatenation-based [25], multiplication-based [26] and attention-based methods [27] In digital pathology, Höhn et al. [28] combined CNN-based histologic WSI analysis with patient metadata to improve skin cancer classification. This approach demonstrated the potential of metadata incorporation to refine diagnostic accuracy and support complex medical imaging tasks.

### 2.4 Deep Learning in Neuropathology

Recently, deep learning has been applied in neuropathology to automate the analysis of histopathological images. Perosa et al. [29] developed CNNs that accurately detect markers of Alzheimer’s disease and cerebral amyloid angiopathy. Koga et al. [30] trained an object detection model to identify five tau lesion types in tau-immunostained digital slide images, enabling the differentiation of tauopathies. Signaevsky et al. [31] applied deep learning techniques to identify the accumulation of abnormal tau in neurofibrillary tangles (NFTs). However, these approaches only focused on pathologically stained images and relied on manually annotated ROIs as the ground-truth labels for training.

Recent studies have demonstrated the potential of MIL in neuropathology. For instance, McKenzie et al. [32] employed an attention-based MIL approach to predict cognitive impairment, identifying key features most associated with cognitive decline. The same group also explored MIL for predicting brain age from neuropathological images, showing its ability to capture age-related features with modest accuracy [33]. Despite these advancements, the application of MIL to CTE and its associated tau pathology remains limited, and the integration of patient metadata with neuropathology images remains largely unaddressed.

## 3 Materials and Methods

### 3.1 Age-informed Attention-based MIL Pipeline

#### 3.1.1 WSI Preprocessing

Our WSI preprocessing pipeline builds upon the CLAM toolbox [20]. The process begins with tissue segmentation using color thresholding at low resolution after stain normalization [34] and median blurring. Morphological closing is then applied to smooth tissue contours and filter out artifacts, followed by contour filtering to exclude areas below a minimum size threshold. Non-overlapping tissue patches of size 224 × 224 pixels are extracted from the segmented tissue regions of WSIs acquired at 20× magnification. For WSIs acquired at 40× magnification, the patch size is adjusted to 429 × 429 pixels to maintain consistent physical dimensions. To analyze the association between grey matter (GM) and white matter (WM) structural changes and AT8 density, we further segmented GM and WM using color thresholding following the initial tissue segmentation step.

#### 3.1.2 Attention-based MIL

As illustrated in Fig. 1, our pipeline employs attention-based MIL (ABMIL) [16]implemented based on the CLAM framework [20]. WSIs are divided into N smaller patches of size 224 × 224 or 429 × 429 pixels. These patches are processed through a pre-trained feature embedding network to generate patch-level representations ***h***_***i***_, where i = 1, 2, …N. These representations serve as inputs to a weakly supervised learning network, which assigns an “attention score” ***α***_***i***_ to each patch, quantifying its relevance for the final classification. We used the gated attention mechanism as described in [16]:

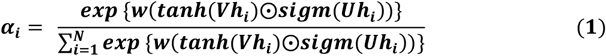

**Fig. 1.**
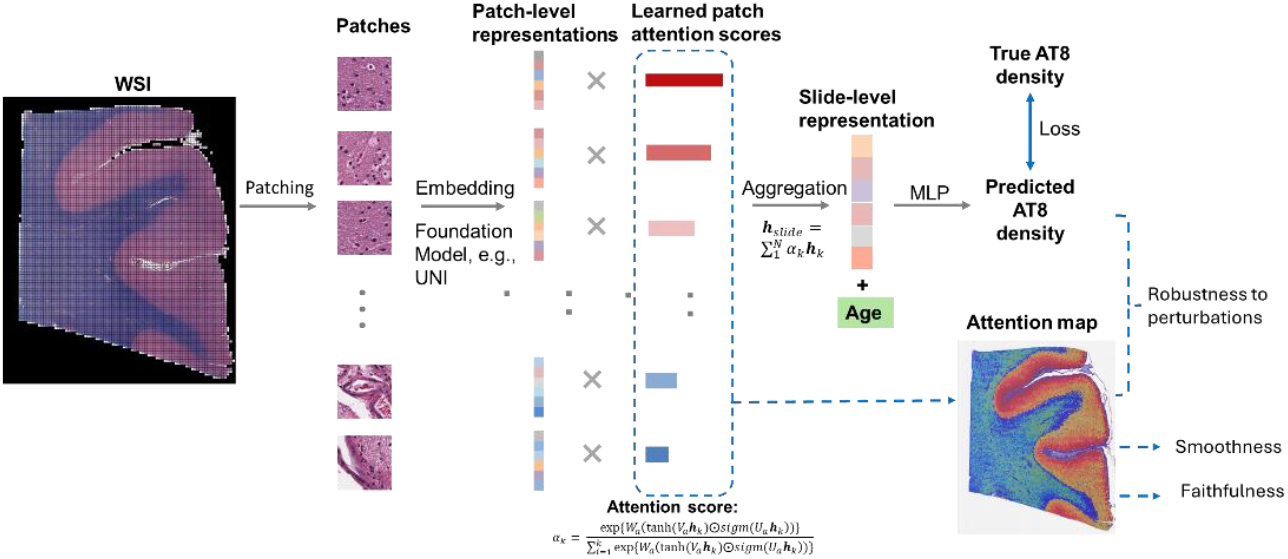
Age-informed Attention-based Multiple-instance Learning Pipeline. WSIs are divided into N smaller patches, which are then processed through a pre-trained feature embedding network (i.e., foundation models like UNI [11]) to generate patch-level representations ***h***_***i***_ . These representations serve as inputs to our weakly supervised learning network, which assigns an “attention score” ***α***_***i***_ to each patch, indicating its relevance in determining the final classification outcome. Subsequently, the patch-level representations are aggregated into a slide-level representation ***h***_***slide***_ using a weighted average, where the weights are the learned attention scores. Based on this slide-level representation, a multi-layer perceptron (MLP) is used to determine the final AT8 density. Quantitative evaluation procedures assesses attention map smoothness, faithfulness, and robustness to perturbations like stain variability and noise for different FMs.

Where **w, V, U** are learnable parameters, ⨀ denotes element-wise multiplication, ***tanh***() is the hyperbolic tangent non-linearity, and ***sigm***() is the sigmoid non-linearity.

The patch-level representations are aggregated into a slide-level representation ***h***_***slide***_ using a weighted average, where the weights are the learned attention scores:

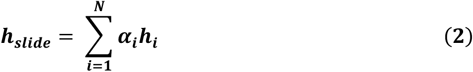

Based on this slide-level representation, a multiplayer perceptron (MLP) with two linear layers and a ReLU activation function between them predicts the AT8 density.

#### 3.1.3 Integration of Patient Age

Higher CTE stages are significantly associated with older age at death [18]. To incorporate patient age into the regression process, ages were normalized to the maximum age value and duplicated 32 times before being concatenated with the slide-level representation ***h***_***slide***_. This multi-modal approach enables the model to leverage both image-based and patient-specific information for improved regression performance.

We also evaluated the performance of different methods for integrating age, including multiplication-based [26]and attention-based [27] methods.

##### Multiplication-based Integration

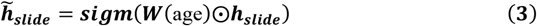

where W is a learnable matrix mapping the age feature to a vector of the same dimension as ***h***_***slide***_.

##### Attention-based Integration

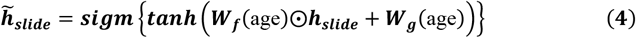

Where ***W***_***f***_ and ***W***_***g***_ are learnable parameters mapping the age feature to a vector of the same dimension as ***h***_***slide***_.

#### 3.1.4 Implementation Details

Our pipeline was implemented in PyTorch. Input embeddings were mapped to an embedding dimension of 512 using a fully connected (FC) layer and then passed into a two-layer gated ABMIL architecture with hidden dimensions of 384. For regularization, dropout with a probability of 0.25 was applied after the first FC layer and within the gated attention layers of the network. The model was trained using the Adam optimizer with a learning rate of 1e-5 and a weight decay of 1e-7. Training was conducted for up to 200 epochs, with early stopping employed based on the validation set loss function.

#### 3.1.5 Evaluation Metrics

The performance of our model was evaluated using the following metrics, including Mean Absolute Error (MAE) that measures the average magnitude of the errors between predicted and actual AT8 densities, and Pearson Correlation Coefficient (PCC) that assesses the linear correlation between predicted and actual AT8 densities, indicating the degree to which the predictions align with true density trends.

### 3.2 Attention Map Analysis

For each WSI, attention maps were generated using the top-performing trained models. In these heatmaps, regions with relatively higher attention are represented in red, while regions with relatively lower attention are represented in blue, with values converted to percentiles across the slide.

To analyze the patches receiving the top-10 highest attention scores as defined by the model, we first converted the RGB images into grayscale and applied Gaussian blurring with a 5 × 5 kernel. The grayscale images were then segmented using multi-Otsu thresholding [35] to isolate nuclei. Morphological openings were applied to smooth boundaries and remove small noise regions from the binarized images. Finally, connected component analysis was used to quantify nuclei features within the patches, including average number of nuclei per patch, average nuclei size, and average nuclei circularity.

### 3.3 Ablation Study

To evaluate the contributions of WM, GM, and patient age to model performance, tissue regions were segmented into WM and GW using color thresholding at low resolution. Separate models were trained and evaluated under various configurations: WM only, GM only, Age only, WM + Age, GM + Age, and WM + GM. The Age-only model was implemented as a support vector machine (SVM) with a nonlinear radial basis function (RBF) kernel.

### 3.4 Evaluation of different FMs as feature extractors

#### 3.4.1 Slide-level Prediction Accuracy

To assess the effectiveness of various pre-trained FMs as feature extractors for neuropathology, we evaluated their performance using predictive metrics. Specifically, MAE and PCC were employed to measure the models’ accuracy in predicting AT8 density. These metrics provided insight into each FM’s suitability for slide-level regression tasks.

#### 3.4.2 Smoothness and Faithfulness of Attention Maps

Based on the assumption that biological tissues typically exhibit smooth transitions, we evaluated the smoothness of the attention maps by quantifying their spatial coherence using the Total Variation (TV) metric, which measures spatial coherence. A lower TV value indicates smoother and more spatially coherent attention maps. The TV of an attention map A was calculated as:

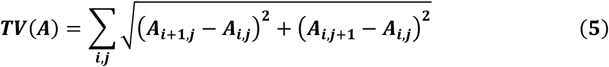

Where ***i, j*** are the index of pixels.

Faithfulness was assessed by masking the top k% and bottom k% of the patches with the highest and lowest attention scores, respectively. The percent change in prediction error (MAE) after masking was used to quantify the model’s dependence on these regions, calculated as:

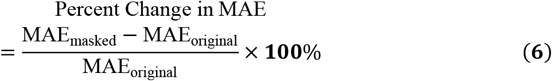

#### 3.4.3 Robustness to Perturbations

Model robustness was evaluated by analyzing performance during inference stage under controlled perturbations. Two types of perturbations were introduced:

##### Random Gaussian noise

Noise following a normal distribution with a mean of 0 and a standard deviation of 0.02 was added to all three color channels (range [0, 1]) to evaluate the model’s robustness to noise.

##### Stain mixing using the REET toolbox [36]

A perturbation parameter ϕ, sampled from a uniform distribution in [-0.05, 0.05], was applied to the color deconvolution matrix R used for transforming RGB space into HED space [37].

The impact of these perturbations was quantified using Percent Change in MAE of the model predictions, as well as the Structural Similarity Index (SSIM) between the original and perturbed attention maps. The SSIM is defined as:

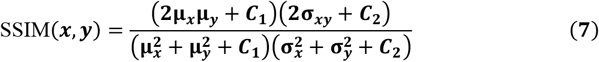

Where ***x*** and ***y*** are the original and perturbed attention map, ***μ***_***x***_ and ***μ***_***y***_ are their average values, 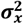 and 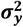 are their variances, and ***σ***_*xy*_ is their covariance. Constants ***C***_**1**_ and ***C***_**2**_ were included to stabilize calculations when denominators are small. The SSIM implementation from the `skimage` package was used.

### 3.5 The Dataset

A detailed description of the whole slide image dataset used in this study is given in Table I. The samples are from deceased individuals with a history of exposure to repetitive head impacts (RHI) from donations to BU Alzheimer’s Disease Research Center (ADRC) brain bank. The CTE staging follows the recent NINDS–NIBIB consensus criteria [4], which includes RHI (patients exposed to RHI without developing CTE), Low CTE (patients with CTE stage I or II), and High CTE (patients with CTE stage III or IV). Cases were excluded from the study if they carried a comorbid neuropathologic diagnosis of Alzheimer’s disease (AD), neocortical Lewy bodies, frontotemporal lobar degeneration (FTLD), or motor neuron disease (MND), based on established neuropathologic criteria for each disease.

**Table 1.** The dataset

| Class | # of patients | Age<br>(mean ± std) | AT8+ cells/mm<sup>2</sup><br>(mean ± std) |
| --- | --- | --- | --- |
| RHI | 137 | 45 ± 22 | 0.528 ± 3.583 |
| Low CTE | 128 | $47 \pm 17$ | $0.358 \pm 0.807$ |
| High CTE | 177 | $69 \pm 22$ | $6.944 \pm 12.703$ |
| All Cases | 438 | $55 \pm 21$ | $3.075 \pm 8.920$ |

Four hundred and thirty-eight total cases met inclusion criteria of having exposure to RHI and not having a comorbid neurodegenerative disease. Formalin fixed paraffin embedded histologic slides were taken from the dorsolateral frontal cortex (Broadman area 6 and 8). AT8 was used to stain for p-tau, and LH&E was used to stain for structural markers. The number of patients for each class is 137, 128, and 177 for RHI, Low CTE, and High CTE, respectively, with each individual contributing one LH&E-stained WSI. Using a GT450 digital slide scanner (Aperio), the WSIs were acquired at either 40× magnification with a resolution of 0.2627 µm per pixel or at 20× magnification with a resolution of 0.5031 µm per pixel. The dataset was divided into training (70%), validation (15%), and test (15%) sets. The AT8 density for each slide was measured by counting the number of AT8-positive cells in the grey matter standardized to the area measured resulting in a density measurement of AT8+ cells /mm^2^. To address the wide range of AT8 density values, we applied a square root transformation to the original AT8 density.

## 4 Results

### 4.1 AT8 Prediction Results

Predicting AT8 density from LH&E images enables the correlation between structural changes in brain tissues and CTE progression. Figure 2 shows the predicted AT8 density versus the ground truth AT8 density for the test set of five folds, respectively. The predicted AT8 densities present a good linear relation with the labels, with PCC ranging from 0.453 to 0.763.

**Fig. 2.**
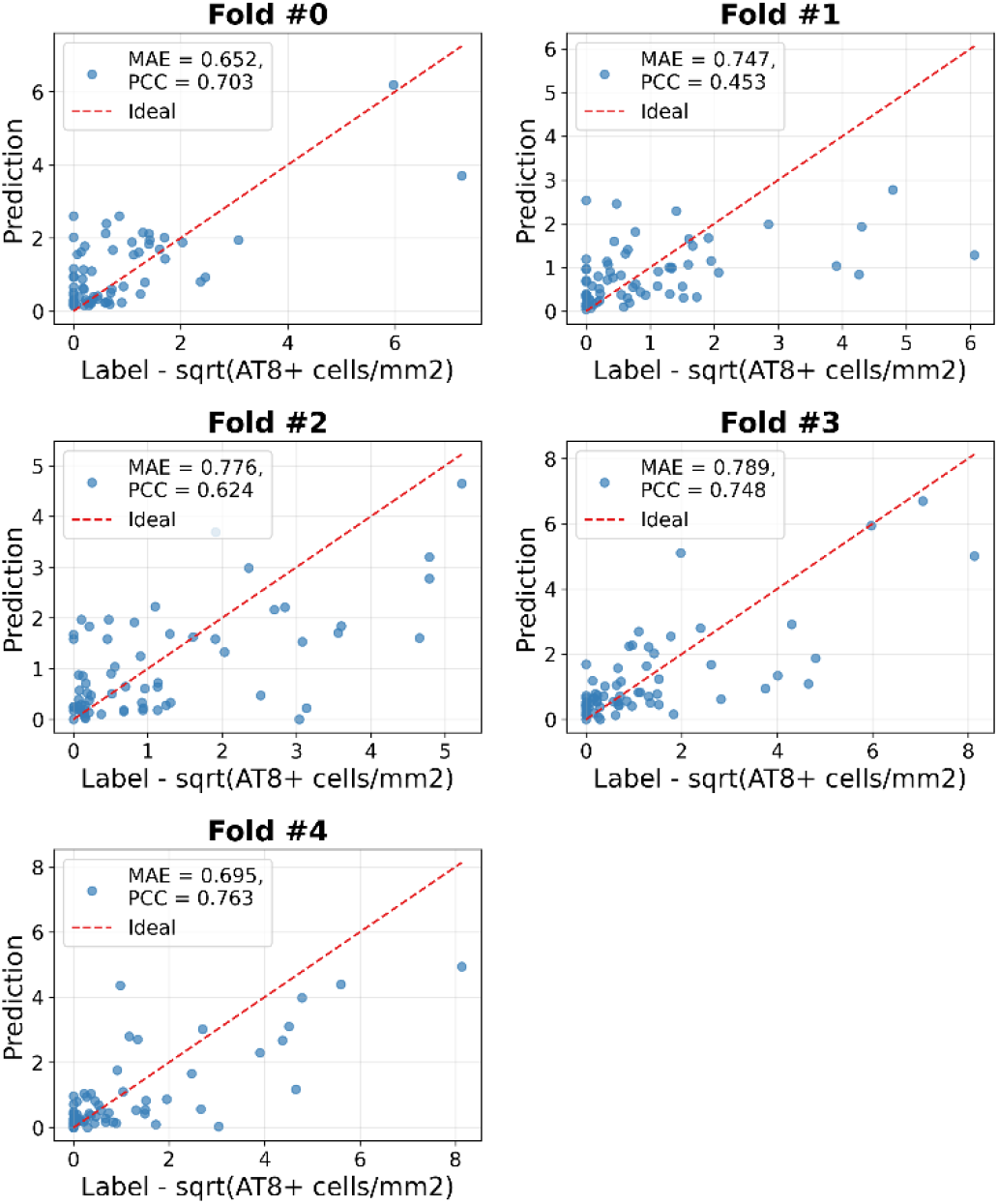
Predicted AT8 density versus the ground truth AT8 density of testing cases for five folds.

### 4.2 Attention maps and interpretation

A critical objective of this part of the study is to identify biologically meaningful regions in LH&E images correlated with known neuropathological changes from AT8 staining. Figure 3 presents examples of LH&E-stained images and their corresponding attention maps generated by our model for representative WSIs. Fold #4 was used in this analysis and all subsequent analyses unless otherwise specified. The attention maps identify ROIs that are most relevant to the model’s regression decision. Notably, the model emphasizes superficial cortical layers (layers I-III), which is consistent with previous findings in CTE research, where NFTs are frequently observed in these regions [6]. This consistency underscores the biological validity of the model’s focus and reinforces its potential as a robust tool for neuropathological investigation.

**Fig. 3.**
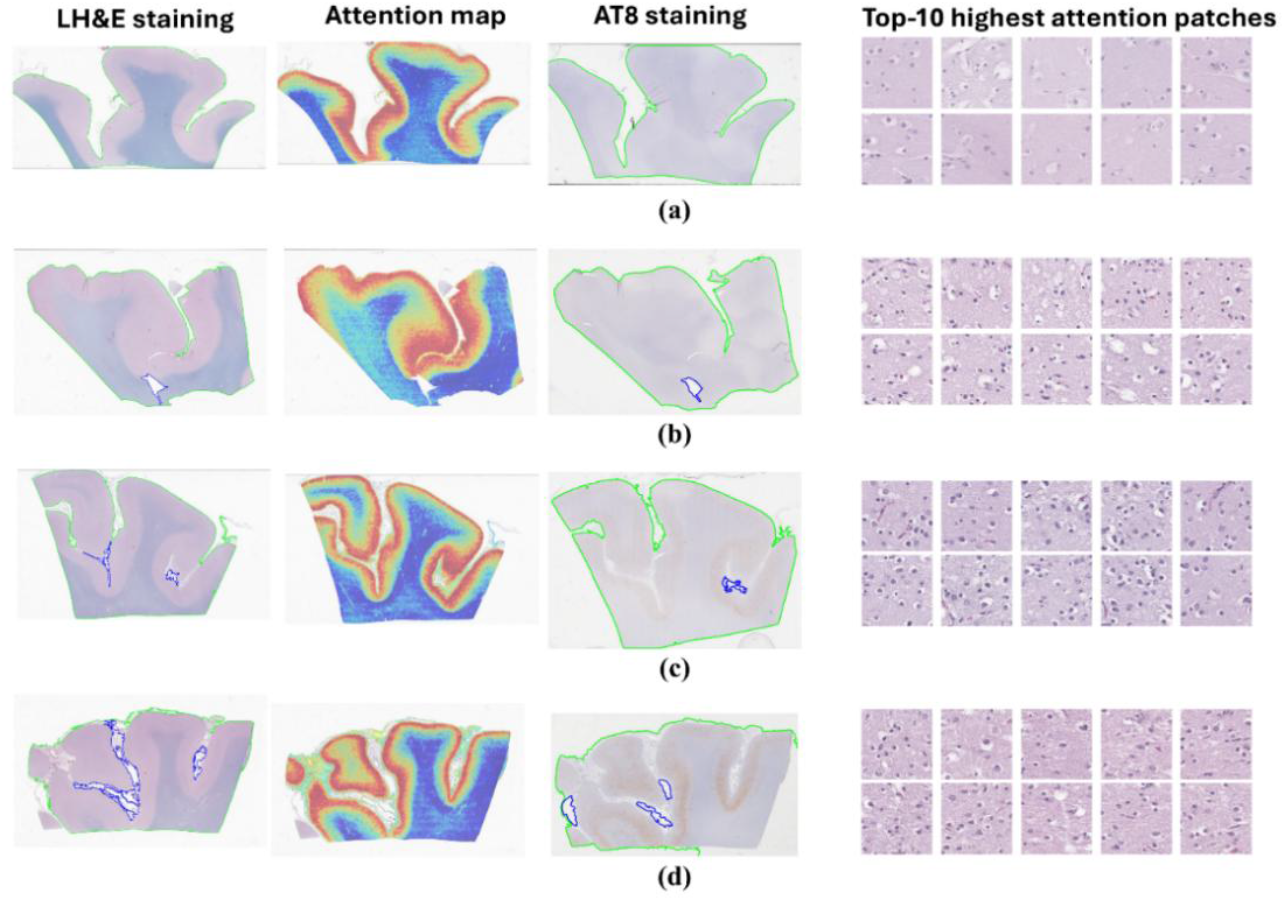
Examples of LH&E staining, corresponding attention maps, corresponding AT8 staining, and top-10 highest attention patches. (a) 0 AT8+ cells/mm^2^ (predicted = 0.01 AT8+ cells/mm^2^) (b) 0.5 AT8+ cells/mm^2^ (predicted = 0.2 AT8+ cells/mm^2^) (c) 31.4 AT8+ cells/mm^2^ (predicted= 19.3 AT8+ cells/mm^2^) (d) 66.1 AT8+ cells/mm^2^ (predicted = 24.4 AT8+ cells/mm^2^).

Figure 4(a) shows one example of the nuclei segmentation from one high-attention patch, and (b)-(d) show averaged number of nuclei in each patch, averaged nuclei sizes, and averaged nuclei circularities the versus the ground truth AT8 density (training, validation, and testing cases are included, and one case was excluded due to failed nuclei segmentation). The cell densities and nuclei sizes show a positive correlation with AT8 density in the top 10 highest-attention patches, while the circularity of the nuclei exhibits a negative correlation. These findings showed specific GM structural alterations with tau aggregation in RHI patients, which is in line with past work [38].

**Fig. 4.**
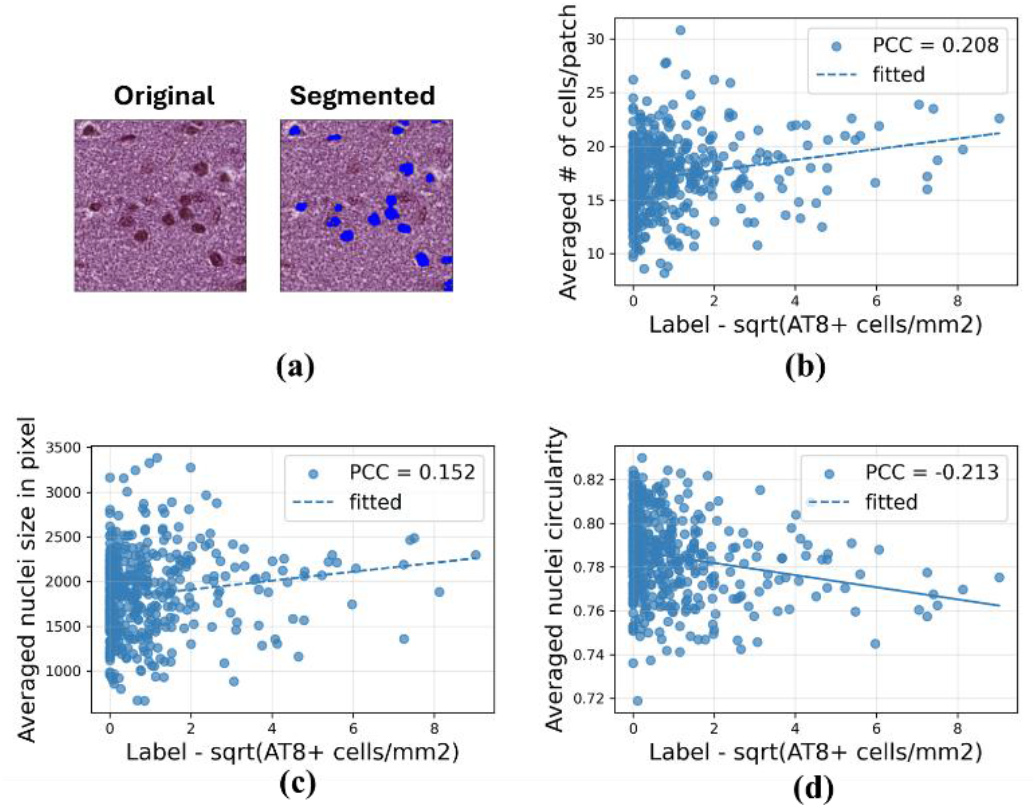
Analysis of top-10 highest attention patches. (a) Example of the nuclei segmentation using multi-Otsu thresholding. (b)-(d) Averaged number of nuclei in each patch, averaged nuclei sizes, and averaged nuclei circularities the versus the ground truth AT8 density.

### 4.3 Ablation study:Evaluating Feature Contributions

To determine the impact of features derived from distinct brain regions (WM and GM) and patient age on the model’s ability to predict p-tau aggregation, Table 2 shows the MAE, and PCC for models trained using image features from different brain regions and the inclusion of the age feature vector. The best performance is achieved by the model using a combination of WM, GM, and age features, with an MAE of 0.734 ± 0.051 and a PCC of 0.658 ± 0.113. The combination of WM and GM features alone also performs well, achieving an MAE of 0.817 ± 0.063 and a PCC of 0.648 ± 0.068. Models using either GM or WM individually yield higher MAE values (0.889 ± 0.079 for GM and 0.921 ± 0.060 for WM) and lower PCC values (0.458 ± 0.099 for GM and 0.461 ± 0.067 for WM). The better performance of GM compared to WM aligns with attention map findings, reinforcing the critical role of GM structural changes in p-tau aggregation.

**Table 2.** Ablation study. The best results are highlighted in bold while the second-best results are marked with an underline. This format is consistent across all subsequent tables.

| Model | MAE<br>(mean ± std) | PCC<br>(mean ± std) |
| --- | --- | --- |
| WM + GM + Age | **$0.734 \pm 0.051$** | **$0.658 \pm 0.113$** |
| WM + Age | $0.889 \pm 0.079$ | $0.458 \pm 0.099$ |
| GM + Age | $0.755 \pm 0.067$ | $0.652 \pm 0.108$ |
| WM + GM | $0.817 \pm 0.063$ | $0.648 \pm 0.068$ |
| WM | $0.921 \pm 0.060$ | $0.461 \pm 0.067$ |
| GM | $0.889 \pm 0.079$ | $0.458 \pm 0.099$ |
| Age (SVM) | $0.781 \pm 0.106$ | $0.515 \pm 0.068$ |

Incorporating age into the model improved predictive performance, particularly for GM. The GM + Age model achieved the second-best performance, with an MAE of 0.755 ± 0.067 and a PCC of 0.652 ± 0.108. Interestingly, the age-only model using a support vector machine (SVM) performed comparably, with an MAE of 0.781 ± 0.106 and a PCC of 0.515 ± 0.068. These results highlight the value of multi-feature integration, particularly the combination of WM, GM, and age features, in achieving superior predictive accuracy.

### 4.4 Evaluation of age integration methods

To assess different strategies for integrating age as a feature [25–27] and determine their impact on model performance when combined with image features, Table 3 presents the MAE and PCC for different methods of integrating age into the model, with WM and GM features included across all experiments. Among the approaches tested, the concatenation-based method achieved the best performance, with an MAE of 0.734 ± 0.051 and a PCC of 0.658 ± 0.113. When comparing all the age-informed models with the base WM + GM model in Table II, our results highlight that both concatenation-based and multiplication-based methods for the inclusion of age feature improves the prediction of AT8 density.

**Table 3.** AT8 prediction results for the WM+GM+Age model with different age integration methods.

| Model | MAE<br>(mean ± std) | PCC<br>(mean ± std) |
| --- | --- | --- |
| Concatenation-based | $0.734 \pm 0.051$ | $0.658 \pm 0.113$ |
| Multiplication-based | $0.795 \pm 0.056$ | $0.602 \pm 0.138$ |
| Attention-based | $0.856 \pm 0.106$ | $0.515 \pm 0.090$ |

### 4.5 Evaluation of different FMs as feature extractors

An important goal of this work is to evaluate how well different FMs perform as a pre-trained feature extractor on neuropathological images with LH&E stains.

#### 4.5.1 AT8 Prediction Results

Table 4 shows the AT8 prediction performance using different feature extractors, including UNI [11], CTransPath [9], Prov-GigaPath [10], and Virchow[15] that are trained on pathology slides, and ResNet [39] which are trained on natural images. The model using UNI achieved the highest PCC and lowest MAE.

**Table 4.** AT8 prediction results for the WM+GM+Age model with different foundation models.

| Model | MAE<br>(mean ± std) | PCC<br>(mean ± std) |
| --- | --- | --- |
| UNI [11] | **0.734** ± 0.051 | **0.658** ± 0.113 |
| CTransPath [9] | 0.819 ± 0.059 | 0.577 ± 0.113 |
| Prov-GigaPath (patch-level) | <u>0.779</u> ± 0.086 | <u>0.622</u> ± 0.071 |
| Virchow [15] | 0.843 ± 0.030 | 0.590 ± 0.090 |
| ResNet [39] | 0.838 ± 0.075 | 0.512 ± 0.060 |

#### 4.5.2 Attention Map Metrics

Figure 5 shows examples of heatmaps from models using different FMs as feature extractors. All attention maps consistently highlighted cortical layers I-III, indicating some level of alignment across models. Visually, the attention maps generated with UNI and ResNet appear smoother than those produced by Prov-GigaPath, CTransPath, and Virchow.

**Fig. 5.**
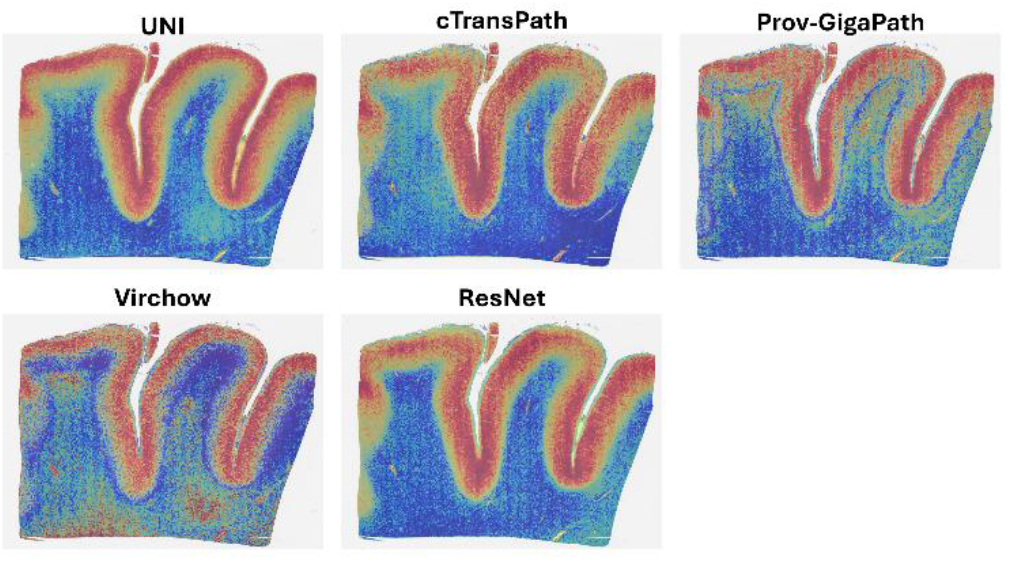
Examples of heatmaps from models using different FMs as feature extractors.

Table 5 presents the smoothness and faithfulness metrics of the attention maps generated by models employing different FMs. Smoothness, measured by TV, indicates that the UNI and ResNet models (with TV values of 8231 and 8224, respectively) produced smoother attention maps compared to Prov-GigaPath (10606), CTransPath (9524), and Virchow (11296). This suggests that UNI and ResNet generate more visually cohesive attention regions.

**Table 5.** Attention map metrics for models using different foundation models.

| Model | Smoothness<br>(Total Variation) | Faithfulness<br>(% MAE change in prediction) |  |
| --- | --- | --- | --- |
|  |  | Top-20%<br>Masking | Bottom-20%<br>Masking |
| UNI | <u>8231</u> | <u>20.93</u> | <u>-0.74</u> |
| CTransPath | 9524 | 0.85 | 0.12 |
| Prov-GigaPath | 10606 | **47.94** | 0.33 |
| Virchow | 11296 | 8.18 | **-1.88** |
| ResNet | **8224** | 7.71 | -0.57 |

For faithfulness, measured by the percentage change in MAE under top-k and bottom-k masking, Prov-GigaPath had the highest MAE change (47.94%) under top-k masking (k = 20%), suggesting that its attention maps are heavily reliant on specific regions for accurate predictions. The UNI model also exhibited a substantial change in MAE under top-k masking (20.93%), indicating that the regions highlighted by its attention maps are highly influential for the model’s predictions. ResNet and Virchow showed a moderate MAE change (7.71% and 8.18%), while in contrast, CTransPath displayed the lowest MAE change (0.85%) under top-k masking, which may indicate a more distributed attention mechanism with less dependency on specific regions. In the bottom-k masking, all models exhibited MAE changes of less than 2%, indicating that these models maintain robustness when the least important regions are masked, with minimal impact on predictive accuracy.

#### 4.5.3 Robustness to Perturbations

Table 6 presents the smoothness and faithfulness metrics of models using different FMs. While the predictions from models using CTransPath, Prov-GigaPath, and ResNet, and Virchow are relatively robust, UNI is sensitive to random Gaussian noise with MAE changes by 20%. All models are relatively robust to stain variation with MAE changes less than 3%. All attention maps remain relatively consistent after adding stain mixing, with average SSIM values exceeding 0.96. However, they exhibit a more significant decline in consistency when adding 2% Gaussian noise, especially Virchow.

**Table 6.**
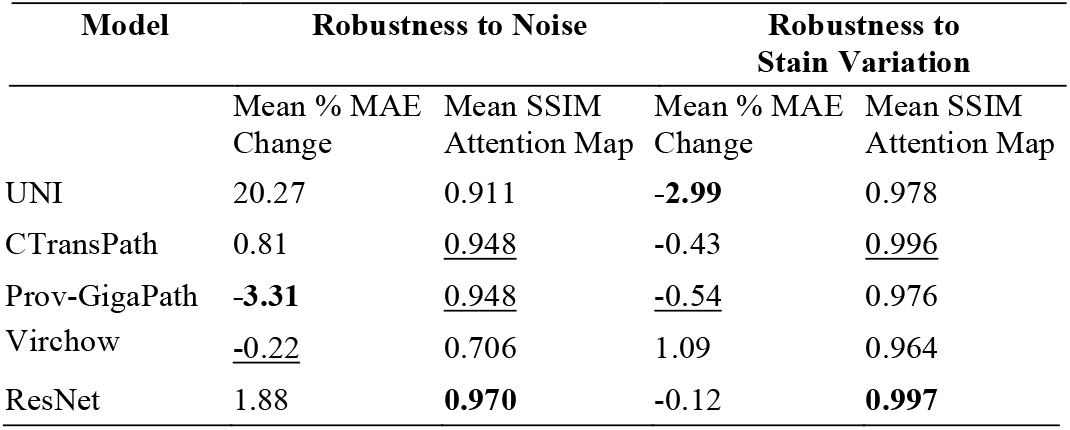
Robustness metrics for models using different foundation models.

## 5 Discussion and Conclusion

In this study, we introduced a novel age-informed, attention-based weakly supervised learning approach for analyzing neuropathological images of brain tissue from individuals with repetitive head impacts (RHI). By focusing on LH&E-stained images and estimating AT8 density, our model provides valuable insights into the correlation between structural changes and tau pathology. The attention maps generated by the model serve as an unbiased indicator of structural changes, offering a new perspective on the neuropathological alterations associated with CTE.

One of the key strengths of our approach is its ability to learn from weak labels, which are more readily available than fully annotated datasets. This is particularly relevant in neuropathology, where obtaining comprehensive annotations can be challenging due to the complexity and variability of the disease. Our method addresses this issue by leveraging the weakly supervised learning framework, which can effectively utilize slide-level labels to train robust models.

The inclusion of age as an informative feature in our model is another significant advancement. Age is a critical factor in the progression of NDs [18, 32] and incorporating it into our model enables more accurate predictions (Table 2) and better contextual understanding of the observed tau pathology. This age-informed approach has the potential to improve the generalizability and applicability of our findings across different patient populations.

Our model highlighted cortical layers I-III, key ROIs in the neuropathological images, correlating well with known patterns of tau pathology in CTE [6]. Previous work has demonstrated that p-tau preferentially aggregates in these layers; however, our findings are the first time to associate specific structural alterations with this region. These findings have important implications for in-life detection of disease. As most NDs can only be definitely diagnosed post-mortem, novel tissue-specific structural alterations, like those observed in this study, may translate to clinical imaging techniques such as MRI, improving our ability to diagnose diseases in living individuals. Overall, these findings underscore the importance of advanced machine learning techniques in augmenting traditional neuropathological methods and enhancing our understanding of complex brain diseases.

A significant contribution of this work is the development of quantitative evaluation procedures for FMs. By introducing metrics such as the smoothness and faithfulness of attention maps, as well as robustness to perturbations including stain variations and random noise, this pipeline establishes a new benchmark for selecting and optimizing FMs for domain-specific tasks. For instance, while the model based on the UNI FM feature extractor [11] achieved the highest prediction accuracy, it performed suboptimally when confronted with noisy data, highlighting the need for robust evaluation measures. These metrics not only enhance the reliability of model outputs but also ensure adaptability to the real-world variability inherent in neuropathological data. Additionally, this framework offers insights for the design of future FMs. For example, CTransPath [9] which incorporates histopathology-specific data augmentation strategies including hue and saturation shifting in HSV space, exhibits greater resilience to staining variability.

However, our study has some limitations. Firstly, the dataset size is relatively small, and the data is derived from a single institution, which may limit the robustness and generalizability of our findings. Secondly, the feature extractors were trained on H&E pathology slides from oncology, potentially limiting their ability to capture features specific to neuropathology LH&E slides. Additionally, while our evaluation procedures for FMs represent a step forward, they remain limited in scope. Incorporating additional perturbations, such as variations in tissue preparation and imaging protocols, could provide a more comprehensive assessment of model robustness. Furthermore, our model’s generalizability to other NDs remains to be investigated. Future work should focus on expanding the dataset to include more diverse staining methods and brain regions and applying our approach to other tauopathies and NDs. Incorporating multi-institutional data and developing standardized scoring criteria could also enhance the model’s applicability and reliability.

In conclusion, this study presents a novel approach to understanding CTE through the use of age-informed, attention-based weakly supervised learning applied to histopathological images. Our methodology effectively estimates AT8 density from LH&E-stained images and generates meaningful attention maps, highlighting structural changes in the brain associated with tau pathology. The robust performance of our model in cross-validation supports its potential for providing new insights into the spatial and structural characteristics of CTE. By leveraging advanced machine learning techniques, we have shown that it is possible to gain a deeper understanding of the neuropathological features of CTE, beyond what is achievable with traditional methods alone. Future studies should aim to extend this framework to other NDs and incorporate a broader range of imaging modalities to deepen our comprehension of brain pathology.

## Abbreviations

ABMIL: Attention-based multiple instance learning
CTE: Chronic Traumatic Encephalopathy
FMs: foundation models
GM: Grey matter
LH&E: Luxol Fast Blue and Hematoxylin & Eosin
MAE: Mean Absolute Error
MIL: Multiple instance learning
NDs: Neurodegenerative disorders
PCC: Pearson Correlation Coefficient
RHI: Repetitive head impacts
ROI: Region of interest
WM: White matter
WSI: Whole-slide image

## Author contributions

SL, AM, JC, and LT: research conception and design. CJ and SL: data curation. SL and MM: methodology and data analysis. SL and JC: results interpretation. SL: manuscript drafting. All authors: manuscript review and editing.

## Funding

Boston University Kilachand Fund and Boston University AD Center (NIA P30AG13846).

## Data availability

The datasets generated and/or analyzed during the current study are not publicly available but are available from the corresponding author on reasonable request.

## Materials availability

Not applicable.

## Code availability

Age-informed ABMIL model, pre-trained model weight, and post-processing codes will be made available on our GitHub page following the publication of this paper.

## Declarations

### Ethics approval and consent to participate

Not applicable. This work does not meet the criteria for human subject research since it only uses postmortem samples.

### Consent for publication

Not applicable.

### Competing interests

The authors declare no competing interests.

## Acknowledgements

We acknowledge funding supports from Boston University Kilachand Fund and Boston University AD Center (NIA P30AG13846).

## Notes

### Competing Interest Statement

The authors have declared no competing interest.

## References

1. Baugh CM, Robbins CA, Stern RA, McKee AC (2014) Current Understanding of Chronic Traumatic Encephalopathy. Curr Treat Options Neurol 16:306. doi:10.1007/s11940-014-0306-5

2. Trojsi F, Christidi F, Migliaccio R, et al (2018) Behavioural and Cognitive Changes in Neurodegenerative Diseases and Brain Injury. Behavioural Neurology 2018:3–5. 10.1155/2018/4935915

3. McKee AC, Cairns NJ, Dickson DW, et al (2016) The first NINDS/NIBIB consensus meeting to define neuropathological criteria for the diagnosis of chronic traumatic encephalopathy. Acta Neuropathol 131:75–86. 10.1007/s00401-015-1515-z

4. Bieniek KF, Cairns NJ, Crary JF, et al (2021) The Second NINDS/NIBIB Consensus Meeting to Define Neuropathological Criteria for the Diagnosis of Chronic Traumatic Encephalopathy. J Neuropathol Exp Neurol 80:210–219. 10.1093/jnen/nlab001

5. Butler MLMD, Dixon E, Stein TD, et al (2022) Tau Pathology in Chronic Traumatic Encephalopathy is Primarily Neuronal. J Neuropathol Exp Neurol 81:773–780. 10.1093/jnen/nlac065

6. McKee AC, Mez J, Abdolmohammadi B, et al (2023) Neuropathologic and Clinical Findings in Young Contact Sport Athletes Exposed to Repetitive Head Impacts. JAMA Neurol 80:1037–1050. 10.1001/jamaneurol.2023.2907

7. Vorontsov E, Bozkurt A, Casson A, et al (2024) A foundation model for clinical-grade computational pathology and rare cancers detection. Nat Med 1–12. 10.1038/s41591-024-03141-0

8. Lu MY, Chen B, Williamson DFK, et al (2024) A visual-language foundation model for computational pathology. Nat Med 30:863–874. 10.1038/s41591-024-02856-4

9. Wang X, Yang S, Zhang J, et al (2022) Transformer-based unsupervised contrastive learning for histopathological image classification. Med Image Anal 81:102559. 10.1016/j.media.2022.102559

10. Xu H, Usuyama N, Bagga J, et al (2024) A whole-slide foundation model for digital pathology from real-world data. Nature 630:181–188. 10.1038/s41586-024-07441-w

11. Chen RJ, Ding T, Lu MY, et al (2024) Towards a general-purpose foundation model for computational pathology. Nat Med 30:850–862. 10.1038/s41591-024-02857-3

12. Zimmermann E, Vorontsov E, Viret J, et al (2024) Virchow 2: Scaling Self-Supervised Mixed Magnification Models in Pathology. 1–19

13. Lu MY, Chen B, Williamson DFK, et al (2024) A visual-language foundation model for computational pathology. Nat Med 30:863–874. 10.1038/s41591-024-02856-4

14. Huang Z, Bianchi F, Yuksekgonul M, et al (2023) A visual–language foundation model for pathology image analysis using medical Twitter. Nat Med 29:2307–2316. 10.1038/s41591-023-02504-3

15. Vorontsov E, Bozkurt A, Casson A, et al (2024) A foundation model for clinical-grade computational pathology and rare cancers detection. Nat Med. 10.1038/s41591-024-03141-0

16. Ilse M, Tomczak JM, Welling M (2018) Attention-based deep multiple instance learning. 35th International Conference on Machine Learning, ICML 2018 5:3376–3391

17. Shao Z, Bian H, Chen Y, et al (2021) TransMIL: Transformer based Correlated Multiple Instance Learning for Whole Slide Image Classification. In: Advances in Neural Information Processing Systems. pp 2136–2147

18. Alosco ML, Cherry JD, Huber BR, et al (2020) Characterizing tau deposition in chronic traumatic encephalopathy (CTE): utility of the McKee CTE staging scheme. Acta Neuropathol 140:495–512. 10.1007/s00401-020-02197-9

19. Quellec G, Cazuguel G, Cochener B, Lamard M (2017) Multiple-Instance Learning for Medical Image and Video Analysis. IEEE Rev Biomed Eng 10:213–234. 10.1109/RBME.2017.2651164

20. Lu MY, Williamson DFK, Chen TY, et al (2021) Data-efficient and weakly supervised computational pathology on whole-slide images. Nat Biomed Eng 5:555–570. 10.1038/s41551-020-00682-w

21. Jaume G, Vaidya A, Chen R, et al (2023) Modeling Dense Multimodal Interactions Between Biological Pathways and Histology for Survival Prediction

22. Bulten W, Kartasalo K, Chen PHC, et al (2022) Artificial intelligence for diagnosis and Gleason grading of prostate cancer: the PANDA challenge. Nat Med 28:154–163. 10.1038/s41591-021-01620-2

23. Zhu J, Liu M, Li X (2022) Progress on deep learning in digital pathology of breast cancer: a narrative review. Gland Surg 11:751–766. 10.21037/gs-22-11

24. Campanella G, Hanna MG, Geneslaw L, et al (2019) Clinical-grade computational pathology using weakly supervised deep learning on whole slide images. Nat Med 25:1301–1309. 10.1038/s41591-019-0508-1

25. Thomas SA (2021) Combining Image Features and Patient Metadata to Enhance Transfer Learning. Proceedings of the Annual International Conference of the IEEE Engineering in Medicine and Biology Society, EMBS 2021-Janua:2660–2663. 10.1109/EMBC46164.2021.9630047

26. Li W, Zhuang J, Wang R, Zhang J (2020) Fusing Metadata and Dermoscopy Images For Skin Disease Diagnosis. 1996–2000

27. Pacheco AGC, Krohling RA (2021) An Attention-Based Mechanism to Combine Images and Metadata in Deep Learning Models Applied to Skin Cancer Classification. IEEE J Biomed Health Inform 25:3554–3563. 10.1109/JBHI.2021.3062002

28. Höhn J, Krieghoff-Henning E, Jutzi TB, et al (2021) Combining CNN-based histologic whole slide image analysis and patient data to improve skin cancer classification. Eur J Cancer 149:94–101. 10.1016/j.ejca.2021.02.032

29. Perosa V, Scherlek AA, Kozberg MG, et al (2021) Deep learning assisted quantitative assessment of histopathological markers of Alzheimer’s disease and cerebral amyloid angiopathy. Acta Neuropathol Commun 9:1–13. 10.1186/s40478-021-01235-1

30. Koga S, Ikeda2 A, Dickson DW (2022) Deep learning-based model for diagnosing Alzheimer s disease and tauopathies. Neuropathology Appl Neurobio 48:

31. Signaevsky M, Prastawa M, Farrell K, et al (2019) Artificial intelligence in neuropathology: deep learning-based assessment of tauopathy. Laboratory Investigation 99:1019–1029. 10.1038/s41374-019-0202-4

32. McKenzie AT, Marx GA, Koenigsberg D, et al (2022) Interpretable deep learning of myelin histopathology in age-related cognitive impairment. Acta Neuropathol Commun 10:131. 10.1186/s40478-022-01425-5

33. Marx GA, Kauffman J, McKenzie AT, et al (2023) Histopathologic brain age estimation via multiple instance learning. Acta Neuropathol 146:785–802. 10.1007/s00401-023-02636-3

34. Reinhard E, Ashikhmin M, Gooch B, Shirley P (2001) Color transfer between images. IEEE Comput Graph Appl 21:34–41. 10.1109/38.946629

35. Liao PS, Chen TS, Chung PC (2001) A fast algorithm for multilevel thresholding. Journal of Information Science and Engineering 17:713–727. 10.1688/JISE.2001.17.5.1

36. Foote A, Asif A, Rajpoot N, Minhas F (2022) REET: Robustness evaluation and enhancement toolbox for computational pathology. Bioinformatics 38:3312–3314. 10.1093/bioinformatics/btac315

37. Ruifrok AC, Johnston DA (2001) Quantification of histochemical staining by color deconvolution. Anal Quant Cytol Histol 23: 23:291–299

38. Nicks R, Shah A, Stathas SA, et al (2024) Neurodegeneration in the cortical sulcus is a feature of chronic traumatic encephalopathy and associated with repetitive head impacts. Acta Neuropathol 148:. 10.1007/s00401-024-02833-8

39. He K, Zhang X, Ren S, Sun J (2016) Deep residual learning for image recognition. Proceedings of the IEEE Computer Society Conference on Computer Vision and Pattern Recognition 2016-Decem:770–778. 10.1109/CVPR.2016.90

